# Redundancy and specificity of type VI secretion *vgrG* loci in antibacterial activity of *Agrobacterium tumefaciens* 1D1609 strain

**DOI:** 10.1101/740209

**Authors:** Mary Nia Santos, Shu-Ting Cho, Chih-Feng Wu, Chun-Ju Chang, Chih-Horng Kuo, Erh-Min Lai

## Abstract

Type VI secretion system (T6SS) is a contractile nanoweapon employed by many Proteobacteria to deliver effectors to kill or inhibit their competitors. One T6SS gene, *vgrG*, encodes a spike protein for effector translocation and is often present as multiple copies in bacterial genomes. Our phylogenomic analyses sampled 48 genomes across diverse Proteobacteria lineages and found ∼70% of them encode multiple VgrGs, yet only four genomes have nearly identical paralogs. Among these four, *Agrobacterium tumefaciens* 1D1609 has the highest *vgrG* redundancy. Compared to *A. tumefaciens* model strain C58 which harbors two *vgrG* genes, 1D1609 encodes four *vgrG* genes (i.e. *vgrGa-d*) with each adjacent to different putative effector genes. Thus, 1D1609 was selected to investigate the functional redundancy and specificity of multiple *vgrG* genes and their associated effectors. Secretion assay of single and multiple *vgrG* deletion mutants demonstrated that these four *vgrG*s are functionally redundant in mediating T6SS secretion. By analyzing various *vgrG* mutants, we found that all except for the divergent *vgrGb* could contribute to 1D1609’s antibacterial activity. Further characterizations of putative effector-immunity gene pairs revealed that *vgrGa*-associated gene 2 (*v2a*) encodes an AHH family nuclease and serves as the major antibacterial toxin. Interestingly, C58’s VgrG2 shares 99% amino acid sequence identity with 1D1609’s VgrGa, VgrGc and VgrGd. This high sequence similarity allows 1D1609 to use an exogenous VgrG delivered from C58 to kill another competing bacterium. Taken together, *Agrobacterium* can use highly similar VgrGs, either produced endogenously or injected from its close relatives, for T6SS-mediated interbacterial competition.

**Author’s Summary:** Selective pressure drives bacteria to develop adaptive strategies, which include competitive and cooperative behaviors. Type VI secretion system (T6SS) is one powerful antibacterial and anti-host nanoweapon employed by many Gram-negative bacteria for growth advantages or pathogenesis. A T6SS-harboring bacterium can encode one to multiple VgrG proteins for delivery of cognate effector(s) but the prevalence and biological significance of having sequence redundant *vgrGs* have not been comprehensively explored. In this study, we investigated the extensiveness of having multicopy *vgrG* genes for effector delivery among diverse Proteobacteria with T6SS. Moreover, a plant pathogenic bacterium *Agrobacterium tumefaciens* strain 1D1609 with highest *vgrG* redundancy was selected for detailed characterization of the roles of multiple VgrGs in T6SS secretion and antibacterial activity. We revealed that the majority of Proteobacterial genomes harbor multiple copies of *vgrG* and the expansion of *vgrG* gene clusters contributed to effector diversity and functional redundancy. Furthermore, the near identical VgrG proteins between 1D1609 and its sibling strain C58 can be exchanged for effector delivery in killing another competing bacterium. Such strategy in using exchangeable effector carriers injected from its isogenic sibling or close relatives during T6SS attacks may be a beneficial strategy for agrobacteria to compete in their ecological niche.

## Introduction

Bacteria have evolved many survival strategies, including pathogenesis, competition, and cooperation, to thrive in diverse and changing environments. One of the weapons that they use is type VI secretion system (T6SS), a machinery that delivers effectors to both eukaryotic and prokaryotic target cells. It is present in approximately 25% of the Gram-negative bacterial genomes sequenced (1).

T6SS has an important role in interbacterial competition and provides competitive advantage in microbial community. It can deliver a variety of effectors such as nuclease, amidase, phospholipase, peptidoglycan hydrolase, muramidase, glycosidase, NADase, and ADP-ribosyltransferase into the target bacterial cell by a contractile mechanism (2–4). Contraction of the sheath-like structure drives the inner tube composed of Hcp tipped by VgrG-PAAR and the associated effectors across bacterial membranes to extracellular milieu or into the target cell. After firing, the structure is immediately disassembled into its individual components, which can be recycled to assemble new machinery for continuous firing (5).

The spike component VgrG is homologous to the gp27/gp5 complex or the tail spike of bacteriophage T4 and assembles into a trimeric complex (6). Current knowledge based on studies from several bacterial systems suggests that VgrG is specifically required for delivery of cognate effector(s) encoded in the same *vgrG* genetic module (7, 8). Furthermore, the C-terminal variable region of VgrG is the molecular determinant conferring specificity of effector delivery by binding to its cognate effector directly or via adaptor/chaperone that interacts with a specific effector (9, 10). T6SS adaptors/chaperones including DUF4123-, DUF1795-, and DUF2169-containing proteins are required for loading a specific effector onto the cognate VgrG for delivery (11).

*A. tumefaciens* is a soil Alphaproteobacterium that infects a broad range of dicotyledonous plants and transfers T-DNA, an oncogenic DNA fragment, to plant’s nuclear genome (12, 13). *A. tumefaciens* strain C58 encodes a single T6SS gene cluster and is equipped with three toxins namely type VI <u>a</u>midase <u>e</u>ffector (Tae), type VI <u>D</u>Nase <u>e</u>ffector 1 and 2 (Tde1 and Tde2), in which its toxin activity can be neutralized by its cognate immunity. Tde is a major antibacterial weapon during *in planta* interbacterial competition and its associated VgrG is specifically required for Tde1/2-dependent bacterial killing (9, 14). A gene which encodes DUF2169 is always found between *vgrG2* and *tde2* orthologs across many Proteobacterial classes (9). In *A. tumefaciens* strain C58, a DUF2169-containing protein encoded upstream of *tde2* is required to stabilize Tde2 and for Tde2-mediated antibacterial activity. For *vgrG1*-associated Tde1, Tap1 (encoded upstream of *tde1*) is the specific chaperone/adaptor interacting with Tde1 prior to loading to VgrG1 for formation of Tde1-Tap1-VgrG-PAAR secretion complex (9).

Aside from the representative *A. tumefaciens* model strain C58, there is little knowledge of T6SS available for other *A. tumefaciens* strains. Our recent comparative analysis of T6SS gene clusters from 11 *A. tumefaciens* strains with complete genome sequences revealed that T6SS is present in all sequenced strains belonging to different genomospecies (15). The *imp* operon (*impA-N*) encoding core structural or regulatory components and the first five genes (*clpV, tai, tae, hcp, vgrG*) encoded in the *hcp* operon are highly conserved but the *vgrG*-associated downstream genes are variable. While all strains only harbor one *vgrG* gene encoded in the main T6SS gene cluster, additional orphan *vgrG* genes not genetically linked to the main T6SS gene cluster were often identified in some strains.

A T6SS-harboring bacterium can encode one to multiple VgrG proteins, in which several of them were demonstrated to be specifically required for delivery of cognate effector(s) encoded in the same *vgrG* genetic module (7, 8). However, the prevalence and biological significance of *vgrG* redundancy has not been tackled. In this study, *A. tumefaciens* 1D1609 which has the highest number of nearly identical *vgrG* genes among all sampled Proteobacteria lineages was chosen to address this question. We investigated the functional redundancy and specificity of multiple VgrG-effector in the effector delivery strategies of T6SS in the context of interbacterial competition. We generated single to multiple in-frame deletion of *vgrG* mutants to characterize the role of paralogous VgrG proteins and its associated effectors in 1D1609. We found that the four *vgrGs* are functionally redundant in mediating Hcp secretion but also exert both redundancy and specificity in mediating effector delivery for interbacterial competition. We also demonstrated that 1D1609 employs a nuclease effector as the major antibacterial toxin. Importantly, we provided experimental evidence that *A. tumefaciens* can use T6SS in the context of interbacterial competition to exchange VgrG as an effector carrier between its close siblings.

## Results

### Majority of T6SS-possessing Proteobacterial genomes harbor multiple *vgrG* genes

To survey the number of *vgrG* genes in T6SS-encoding bacterial genomes, 48 representative Proteobacteria harboring T6SS gene cluster(s) were selected for phylogenetic analysis. The result revealed that 33 of these genomes (∼70%) encode multiple *vgrG* genes. The *vgrG* copy numbers have no strong tie to the phylogenetic placement of individual genomes (Fig 1), suggesting that the copy number evolution is highly dynamic, even at within-genus level (e.g. *Agrobacterium* in Alphaproteobacteria or *Pseudomonas* in Gammaproteobacteria). The *vgrG* gene tree (Fig S1) is not congruent with the species tree (Fig 1) and the patterns suggest that horizontal gene transfers across different classes are not rare events. Nonetheless, the species with multiple VgrG homologs mostly (∼64%) form a monophyletic clade at family levels. All sampled Rhizobiaceae lineages except for *Sinorhizobium fredii* form a monophyletic clade and has only one main T6SS gene cluster (Fig S1) (15, 16). This pattern suggests that gene duplication could be the major driving force for copy number increase within families.

**Fig 1.**
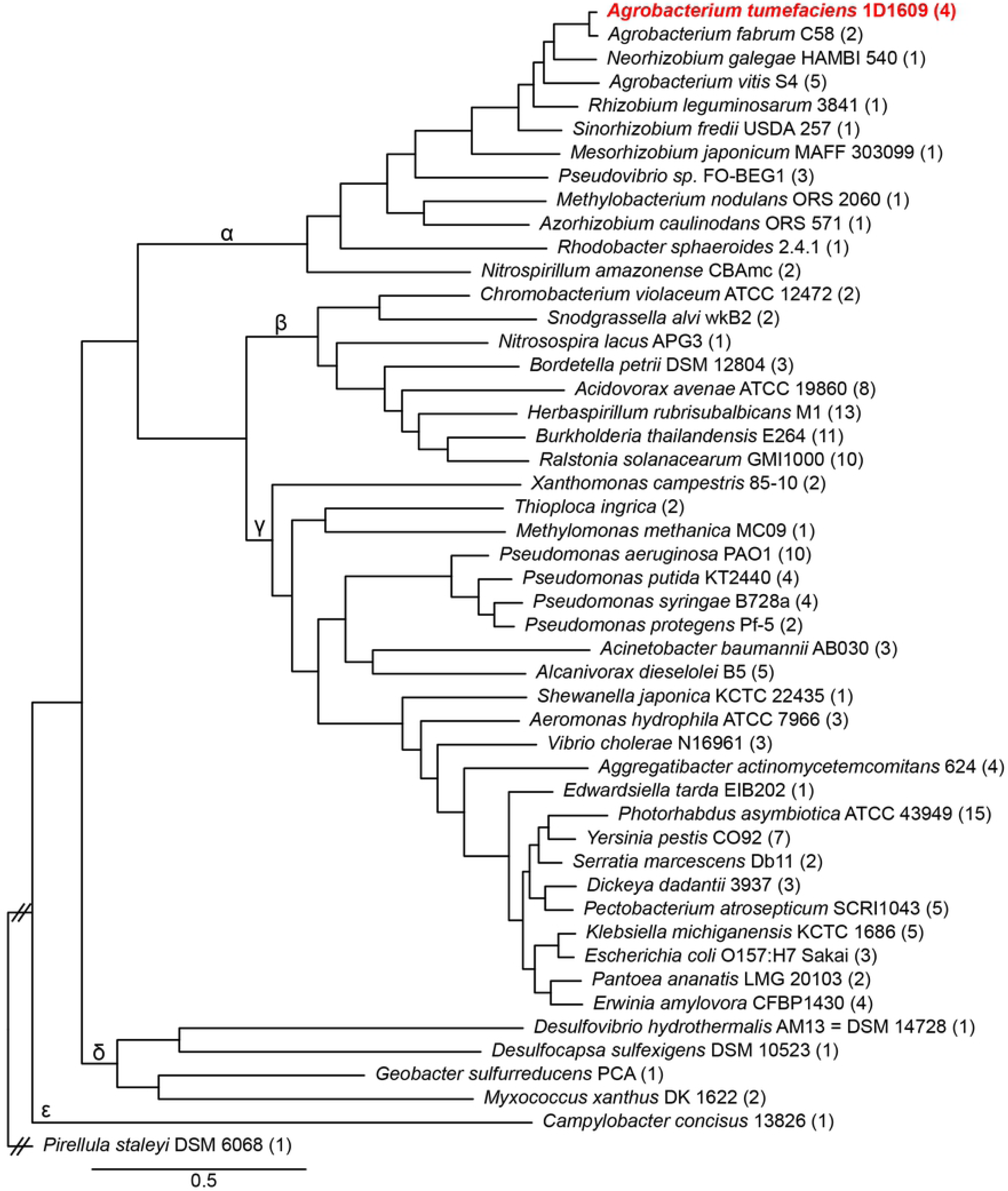
Bayesian phylogeny of representative Proteobacteria based on 120 shared single-copy genes. The labels on the branches indicate the five classes within this phylum: Alphaproteobacteria (α), Betaproteobacteria (β), Deltaproteobacteria (δ), Epsilonproteobacteria (ε) and Gammaproteobacteria (γ). The numbers in parentheses after strain names indicate the *vgrG* copy numbers.

We further asked, when there are multiple *vgrG* homologs in a genome, how often can we find highly similar genes that are likely to be functionally redundant. Using protein sequence identity ≥ 95% as the cut-off, only four genomes met the criteria (Table S1). These include Alphaproteobacterium *A. tumefaciens* strain 1D1609 (three out of four with 99% identity, the remaining one is 93-94% identical), Betaproteobacterium *Acidovorax avenae subsp. avenae* ATCC 19860 (two out of eight with 99% identity), Gammaproteobacterium *Aggregatibacter actinomycetemcomitans* strain 624 (four copies belonging to two types: within-type = 99-100% identity, between-type = 90% identity), and Gammaproteobacterium *Dickeya dadantii* strain 3937 (two out of three with 97% identity, the remaining one is 78% identical). Among them 1D1609 has the highest redundancy.

### 1D1609 has four *vgrG* genes with each genetically linked to specific putative effector gene(s)

1D1609 genome encodes one main T6SS gene cluster including *vgrGa* (At1D1609_RS23245) that is encoded in *hcp* operon within the T6SS main gene cluster. The other three *vgrGs* - *vgrGb* (At1D1609_RS26290)*, vgrGc* (At1D1609_RS22460) and *vgrGd* (At1D1609_RS18895) are encoded in different loci elsewhere (Fig 2). All four *vgrG* genes are genetically linked with distinct potential effector gene(s) and associated genes predicted to function as adaptor/chaperone or immunity (15). The four *vgrG* genetic modules consist of the first three genes with conserved domain and same gene orders encoding VgrG-DUF2169-DUF4150. The *vgrGa-, vgrGc-,* and *vgrGd*-associated genes (*v2a, v2c, v2d*) encode N-terminal PAAR-like DUF4150 domain followed by C-terminal putative effector domain. The *vgrGb*- linked *v2b* does not encode obvious effector domain and instead followed by downstream genes encoding putative effector domain, as predicted by Phyre or NCBI CDD search and BLASTP (17–19) (Table S2). The *vgrGc*-associated putative effector gene *v2c* does not encode any conserved domain, while the one (*v2d) associated* with *vgrGd* encodes a putative GH25 muramidase domain. Genes encoding DUF2169 domain are commonly found downstream of *vgrG* and upstream of DUF4150-containing effector genes. This gene order in *vgrG* genetic module is highly conserved in the *vgrG2* locus of C58, in which DUF2169-containing protein (Atu3641) may function as adaptor/chaperone of Tde2 due to its requirement for Tde2 stability and Tde2-dependent bacterial killing (9). Thus, DUF2169-containing V1 protein is likely to function as a chaperone/adaptor required for delivery of cognate effectors in conjunction with associated VgrG.

**Fig 2.**
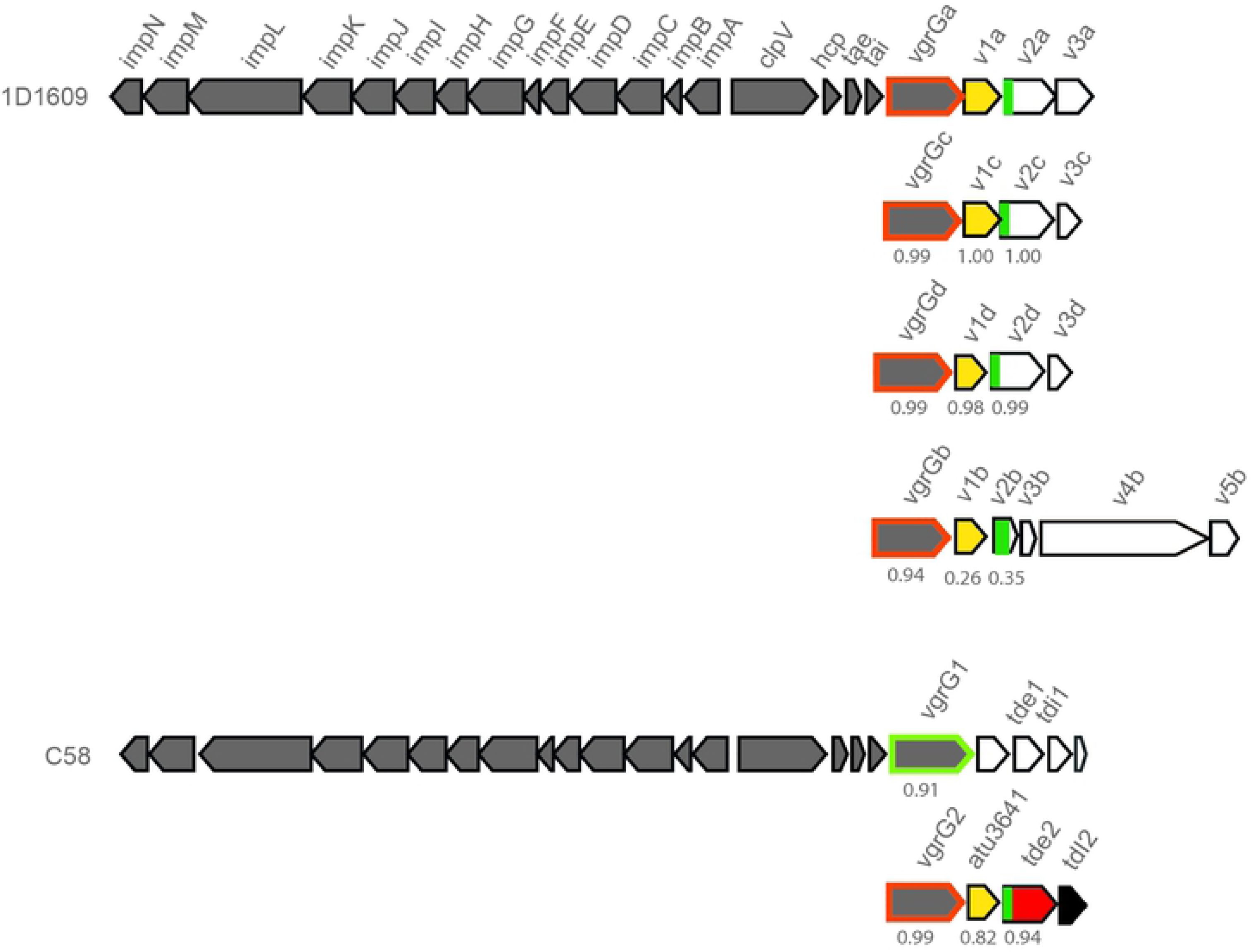
1D1609 encodes four *vgrG* genes, each is genetically linked with different potential effector gene(s). Genetic organization of *A. tumefaciens* 1D1609 and C58 T6SS gene cluster and *vgrG* genetic modules. C58 *vgrG1* is outlined with green and C58 *vgrG2* and 1D1609’s four *vgrG* genes are outlined in orange. The gene encoding DUF2169 domain is highlighted in yellow and DUF4150 (PAAR-like) domain is shown in dark green. The *vgrG-associated genes* (*v*) are numbered according to their position from *vgrG*. The percentage identity, which corresponds to Table S3 is indicated below the gene cluster.

Unlike VgrG1 and VgrG2 which share 92% identity and are highly divergent at their C-terminus (9), we found that the three VgrG proteins – VgrGa, VgrGc and VgrGd in 1D1609 share 99% overall amino acid sequence identity (and 100% identity in the C-terminal region). These are also highly similar with VgrG2 of C58 (Fig 2, Table S3). The VgrGb shares 93-94% overall identity to the other three VgrGs and has a highly divergent C-terminal region (Fig S2, Table S3). Consistent with the VgrG comparisons, the DUF2169 and DUF4150 proteins with *vgrGa*/*c*/*d* also share 98-100% and 99-100% amino acid sequence identity, respectively (Table S3). In contrast, VgrGb-associated DUF2169 and DUF4150 share only 26% and 35% sequence identities to those encoded in the other three *vgrG* loci respectively. The four predicted effector domains fused to DUF4150 or Rhs do not share high sequence similarity to each other or to known effectors, which could equip 1D1609 with diverse effector activities to fight with a wide range of competing bacteria. Homologs of *vgrGa*-, *vgrGb*-, *vgrGc*-, and *vgrGd*-associated effector genes are widely found in species belonging to Rhizobiales, suggesting that these are common effectors in this group.

### Four VgrG proteins are functionally redundant in mediating Hcp secretion

In C58, VgrG1 and VgrG2 are functionally redundant in mediating Hcp secretion but specifically required for delivery of the cognate effectors Tde1 and Tde2, respectively (9, 20). Each of in-frame deletion mutants of *vgrGa*, *vgrGb*, *vgrGc* and *vgrGd* were generated to determine their functions in mediating Hcp secretion, a hallmark of a functional T6SS assembly. The *ΔtssL* mutant, with deletion of gene encoding the essential T6SS membrane component TssL, is used as a negative control. The secretion assay showed that all single *vgrG* deletion mutants remain active in Hcp secretion, suggesting that none of these VgrG proteins are essential for assembly of a functional T6SS (Fig 3A). We are able to detect secretion of VgrG proteins from 1D1609 using polyclonal antibody against VgrG1 protein of C58, which can recognize all four VgrG proteins produced in 1D1609 and VgrG2 of C58. The Hcp and VgrG proteins are secreted in a T6SS-dependent manner because no Hcp and VgrG signals are detected in secretion fraction of *ΔtssL* and RpoA (RNA polymerase α-subunit), a non-secreted protein, is not detectable in any secretion fraction. Hcp secretion remains active in double and triple *vgrG* deletion mutants but is only completely abolished when all four *vgrG* genes was deleted (Fig 3B). The overexpression of each of all four *vgrG* on a plasmid in the quadruple *vgrG* mutant restores Hcp secretion (Fig 3C). Furthermore, trans complementation of each 1D1609 *vgrG* constitutively expressed on a plasmid can also restore Hcp secretion in C58 double *vgrG* deletion mutant (Δ*vgrG*1,2) (Fig 3D). This is consistent with previous study that either VgrG1 or VgrG2 is sufficient in mediating the secretion of Hcp in C58 (20). Indeed, all triple *vgrG* mutants wherein only one VgrG is present remains capable of mediating Hcp and VgrG secretion (Fig 3E). In Δ*vgrGabd* mutant, wherein only VgrGc is produced, Hcp secretion level is at lesser amount as compared to WT. The transcript expression level of *vgrGc* is also low compared to the other *vgrG*s (Haryono et al., 2019). Altogether, our comprehensive secretion assays demonstrated that the four VgrGs are functionally redundant in mediating Hcp secretion. Single *vgrG* allele is sufficient for assembly of a functional T6SS although the degree of assembly efficiency may not be the same due to expression level or sequence divergence.

**Fig 3.**
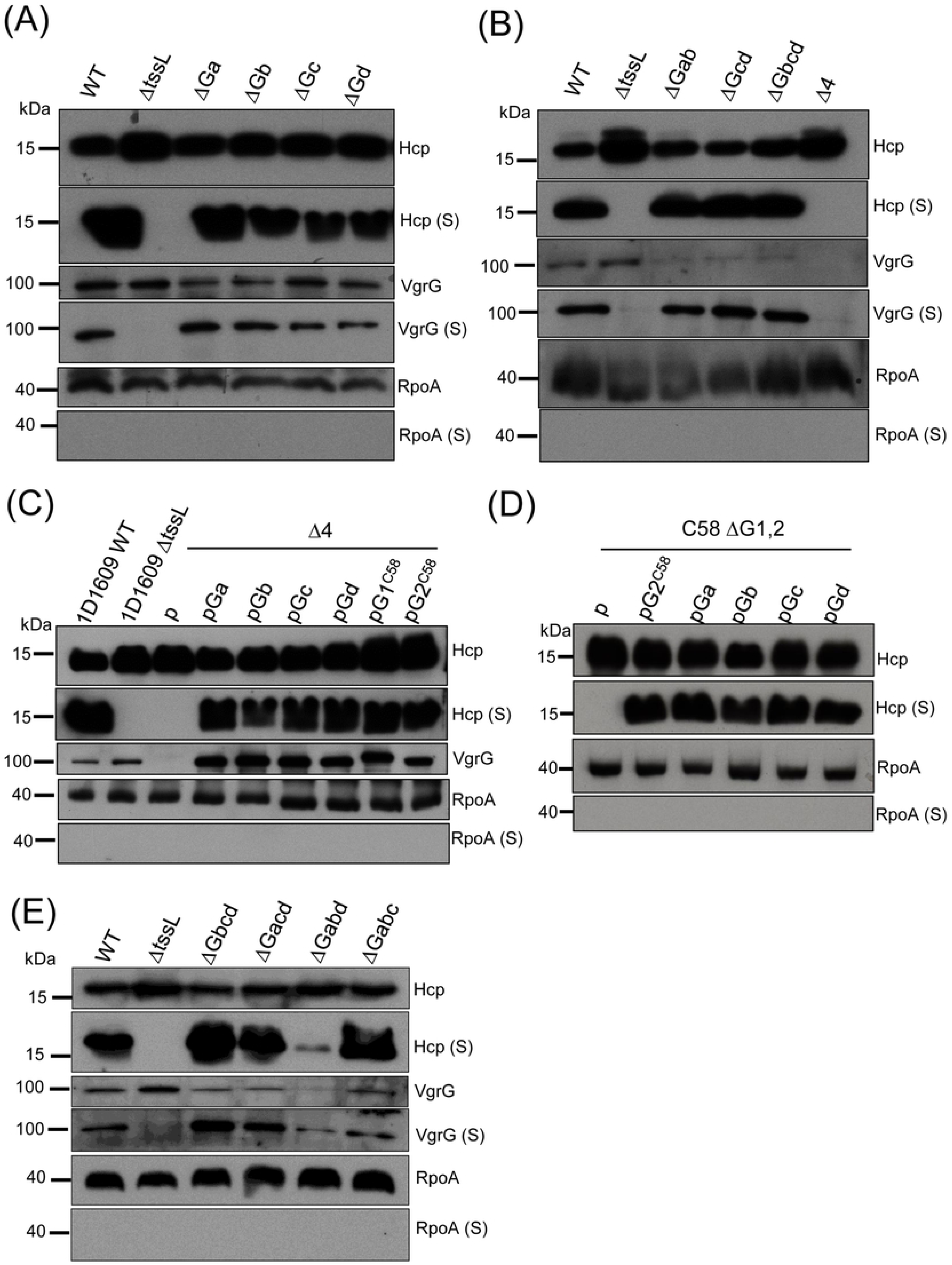
The four VgrG proteins of 1D1609 are functionally redundant for Hcp secretion. Western blot analysis of cellular and secreted (S) proteins from single *vgrG* deletion mutants **(A)**, multiple *vgrG* deletion mutants **(B)**, overexpression of VgrG in 1D1609 quadruple *vgrG* mutant (Δ4) **(C),** and in C58 double *vgrG* deletion mutant (ΔG1,2) **(D),** and triple *vgrG* deletion mutants **(E)**. Various *A. tumefaciens* strains were cultured in LB (A, B and E) or AK (C and E) broth at 25°C and cellular and secreted (S) fractions were subjected to western blotting using anti-Hcp, anti-VgrG, and RpoA as indicated. The T6SS inactive mutant Δ*tssL* is a negative control and RpoA serves as a non-secreted protein control. Protein markers (in kDa) are indicated on the left. Similar results were obtained from two to four independent experiments.

### T6SS-dependent antibacterial activity against *Escherichia coli* is mainly contributed by *vgrGa* and *vgrGd* alleles

We previously showed that all tested *A. tumefaciens* strains including 1D1609 exhibit T6SS-mediated antibacterial activity against T6SS-negative *E. coli* (15). Here, we further investigated the role of the four *vgrG* genes in interbacterial competition against *E. coli*. As a positive control, ∼10x reduction of surviving *E. coli* target cells was observed when co-cultured with 1D1609 WT as compared to that of *ΔtssL.* However, each of single *vgrG* deletion mutants remains to have similar antibacterial activity to WT (Fig 4A). Further antibacterial activity assay of double, triple and quadruple *vgrG* mutants showed that the antibacterial activity of 1D1609 is completely lost in *ΔGacd, ΔGabd*, or *ΔGabcd* while *ΔGbcd* and *ΔGabc* triple *vgrG* mutants remain similar antibacterial activity to that of WT 1D1609 (Fig 4B, C). These results suggest that *vgrGa* and *vgrGd* alone is sufficient to contribute to the antibacterial activity of 1D1609. In contrast, *vgrGb* or *vgrGc* alone cannot exert any detectable antibacterial activity even though both alleles alone (*ΔGacd, ΔGabd*) are sufficient in mediating Hcp and VgrG secretion (Fig. 3E). However, when each of single *vgrG* is constitutively overexpressed on a plasmid in quadruple *vgrG* mutant, *vgrGa, vgrGc*, and *vgrGd* but not *vgrGb* is able to partially restore antibacterial activity (Fig 4D). These results strongly suggest that endogenous *vgrGa* and *vgrGd* indeed play important roles for antibacterial activity while *vgrGb* plays no role in antibacterial activity to *E. coli.* The data that endogenous *vgrGc* plays no role but can exert its function in antibacterial activity when overexpressed is consistent with its low endogenous expression and high amino acid sequence identity to VgrGa and VgrGd.

**Fig 4.**
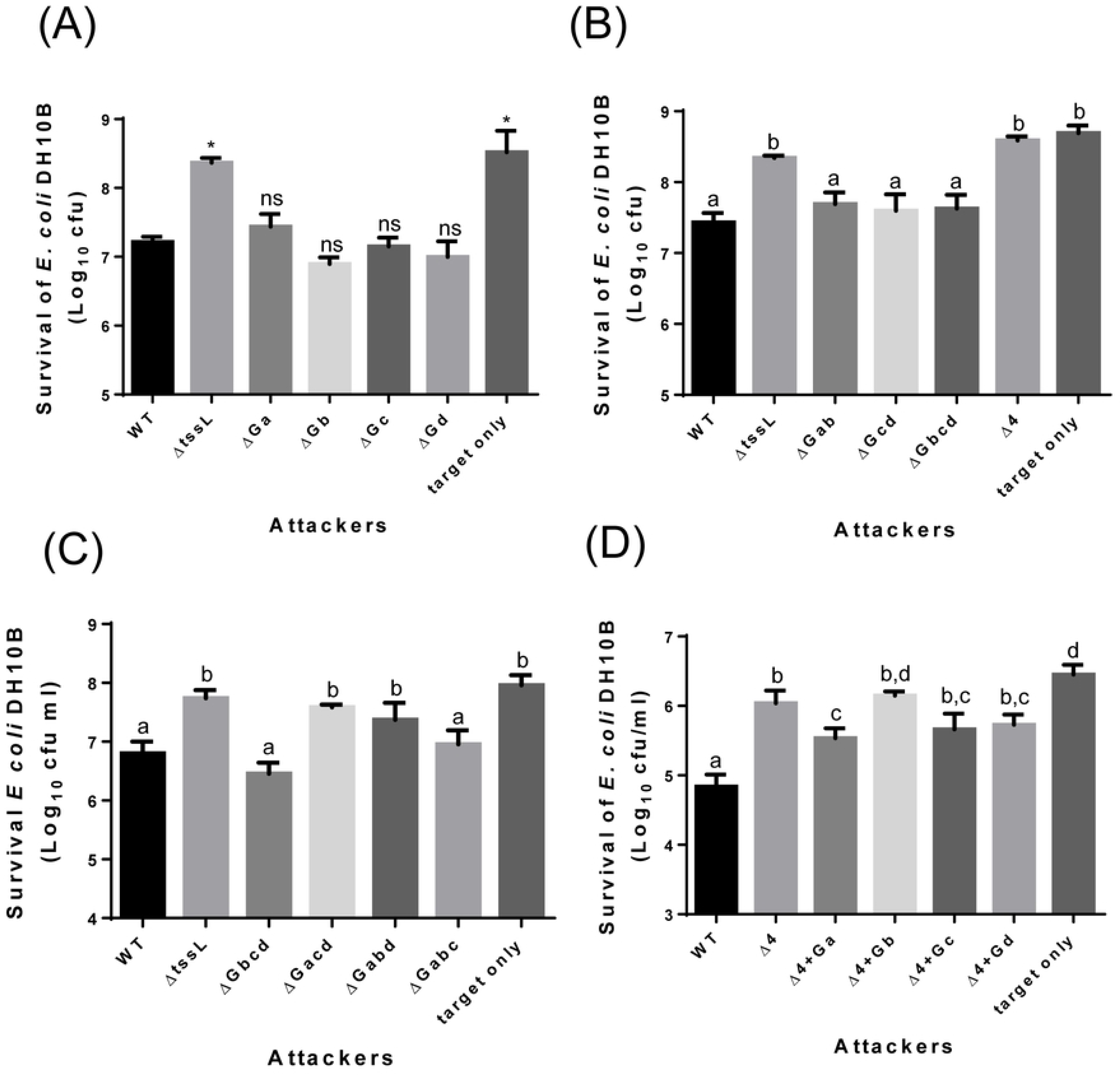
*vgrGa* and *vgrGd* contributes to antibacterial activity of 1D1609. Each of various *A. tumefaciens* 1D1609 strains (indicated in the x axis) was mixed with *E. coli* DH10B cells expressing pRL662 to confer gentamicin resistance at 30:1 ratio in LB (A and B) or AK (C and D) media for 18 hr. The survival of *E. coli* cells was quantified as cfu as shown on the y axis. **(A)** Antibacterial activity of single *vgrG* deletion mutants. Data are mean ± SEM of three biological replicates, similar results were obtained from three independent experiments. Two-tailed Student’s t-test was used for statistical analyses. * p < 0.01. **(B)** Antibacterial activity in multiple *vgrG* deletion mutants. **(C)** Antibacterial activity using triple *vgrG* mutants. **(D)** Single *vgrG* expressed *in trans* using pRLBla in quadruple *vgrG* mutant (Δ4). Data are mean ± SEM of three biological replicates (B) or four biological replicates from two independent experiments (C and D) and letters above the bar indicate significant difference (P < 0.05) determined by Tukey’s HSD test.

### The *vgrGa*- and *vgrGd*-associated EI pairs are responsible for antibacterial activity of 1D1609 and V2a is a nuclease of HNH/ENDO VII superfamily

We also generated deletion mutants of each putative effector-immunity (EI) gene pairs individually and mutants with deletions in multiple putative EI pairs. Due to failure in generating the *vgrGc*-associated *v2c-v3c* EI pair deletion mutant after several attempts, we instead generated the deletion of the whole *vgrGc* cluster to assay the antibacterial effect of *vgrGc*-associated effector. Deletion of *vgrGa*-associated EI pair partially compromised antibacterial activity against *E. coli* while deletion of *vgrGb*-associated EI pair or *vgrGc* cluster/associated toxin has no effect in killing *E. coli*, as compared to WT and *ΔtssL* (Fig 5A). Importantly, any mutant minimally deleting both *vgrGa*- and *vgrGd*-associated EI pairs *(Δ2TIad*, *Δ3TIabd*, or *Δ4TIabcd*) completely abolished the antibacterial activity against *E. coli* (Fig 5B). These results suggest that *vgrGa*- and *vgrGd*-associated EI pairs contribute to antibacterial activity to *E. coli*.

**Fig 5.**
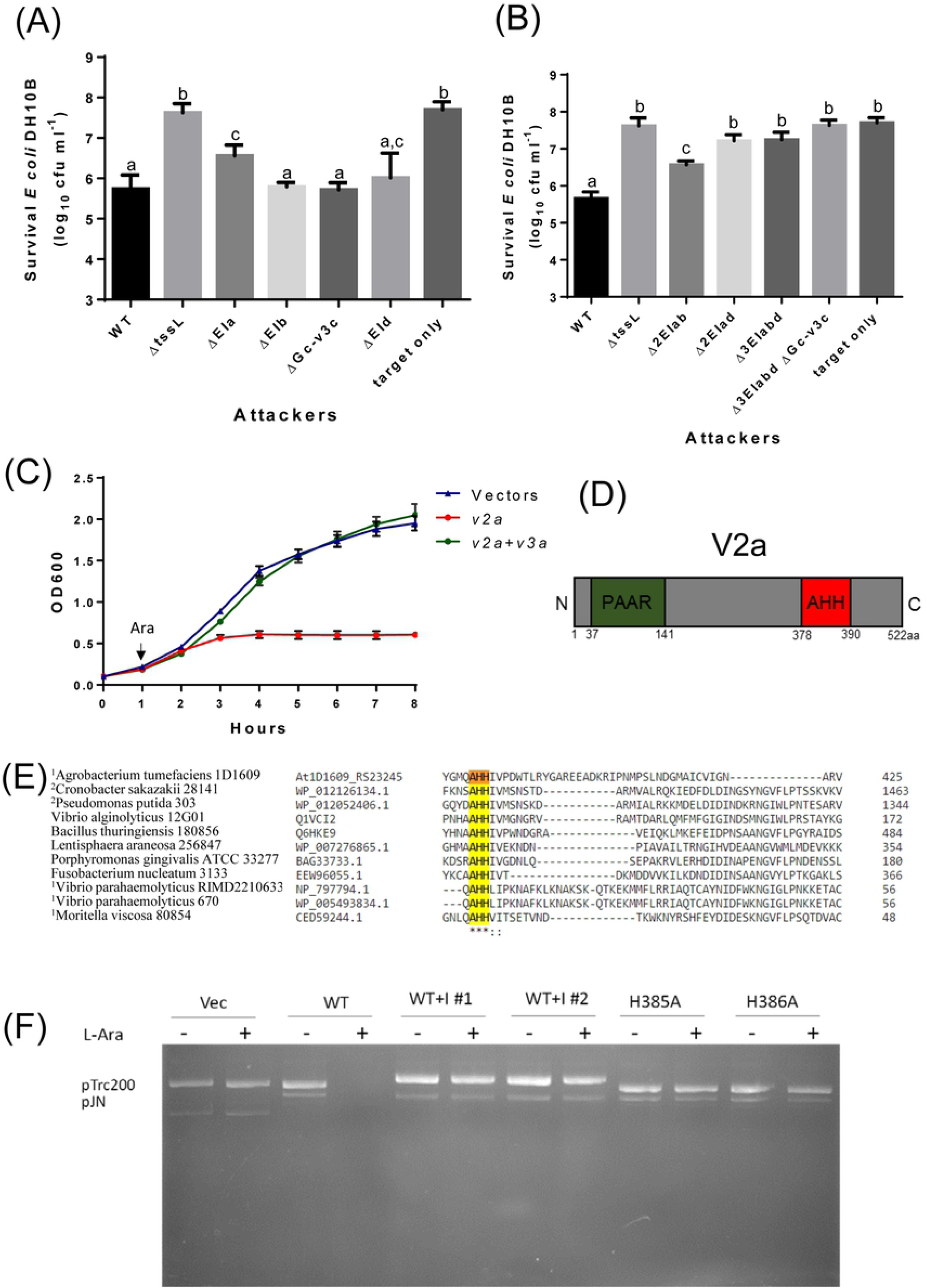
*vgrGa*- and *vgrGd*-associated EI pairs are responsible for antibacterial activity and V2a is a nuclease. *A. tumefaciens* 1D1609 strains (indicated in the x axis) was mixed with *E. coli* DH10B cells expressing pRL662 to confer Gentamicin resistance at 30:1 ratio in LB. The survival of *E. coli* cells was quantified as cfu as shown on the y axis when co-cultured with single **(A)** and multiple **(B)** EI pair deletion mutant. Data are mean ± SEM of four biological repeats from two independent experiments and letters above the bar indicate significant difference (P < 0.05) determined by Tukey’s HSD test. **(C)** *E. coli* growth inhibition analysis in *vgrGa*-associated EI pair. *E. coli* cultures were induced with 1 mM IPTG at 0 hr for the expression of putative immunity protein expressed on pTrc200 plasmid followed by L-arabinose (Ara) induction at 1 hr to induce putative toxin gene from pJN105 plasmid. Cell growth was recorded every hour at OD_600_. Empty vectors were used as controls. Data are mean ± SEM of three independent experiments. **(D)** Schematic diagram showing N-terminal PAAR region and C-terminal AHH nuclease domain of *vgrGa*-associated effector V2a. **(E)** Partial sequence alignment of representative nuclease family of HNH/ENDO VII superfamily with conserved AHH motif, modified from NCBI CDD sequence alignment of nuclease family of the HNH/ENDO VII superfamily with conserved AHH (pfam14412). The strain name and locus tag/accession number are on the left, and the conserved amino acid residues are indicated as * (identical) and : (similar) respectively. AHH motif is highlighted in yellow with the two His targeted for mutagenesis. AHH domain linked to DUF4150 domain^1^ or Rhs^2^ is indicted. **(F)** Nuclease activity assay. *E. coli* cells with pTrc200 and pJN105 plasmid (Vec) or derivatives expressing *v2a* (WT), WT with immunity protein (WT + I; #1 and #2 are two independent constructs) and catalytic site mutants (H385A, H386A) were induced with (+) or without (-) L-arabinose (L-Ara) for 2 hr. Plasmid DNA was extracted and the degradation pattern was observed in agarose gel.

Since single deletion of *vgrGa*-associated EI pair (*ΔEIa*) or in combination with additional EI-pair/s showed reduced or abolished antibacterial activity (Fig 5A and B), we considered *vgrGa*-associated effector V2a as the primary toxin and characterized its function. The putative effector gene *v2a* is expressed by an arabinose inducible promoter on plasmid pJN105 and its putative cognate immunity gene *v3a* was expressed constitutively on plasmid pTrc200 in *E. coli*. The results showed that the bacterial growth is inhibited when *v2a* is induced as compared to the vector control whereas co-expression with *v3a* restores the growth (Fig 5C), indicating their toxin-immunity relationship.

V2a is 522-aa in length and contains a N-terminal PAAR-like DUF4150 domain and a C-terminal nuclease domain belonging to HNH/ENDOVII superfamily (pfam14412) of the treble cleft fold (18) (Fig. 5D). This nuclease family has a conserved motif AHH, which is predicted toxin module found in bacterial polymorphic toxin systems. In the AHH motif, the first His forms one of the catalytic metal-chelating ligands and the second His contributes to the active site that directs the water for phosphoester hydrolysis (21). Partial sequence alignment of representative nuclease family of HNH/ENDO VII superfamily with conserved AHH motif is shown in Fig 5E.

To determine its nuclease activity, we examined the degradation of the plasmid DNA extracted from *E. coli* cells after *v2a* is induced by arabinose for 2 hr. We showed that both pTrc200 and pJN-derived plasmids were not detectable from *v2a*-containing cells induced by arabinose, in contrast to the presence of both plasmids in vector control or *v2a*-containing cells not treated with arabinose (Fig. 5F). The plasmid degradation is indeed caused by nuclease activity of V2a because the plasmid degradation is no longer detectable when the cells expressing the catalytic site (H385A and H386A) mutant or co-expression of *v3a* (Fig. 5F). Taken together, we demonstrated that V2a is a T6SS effector that requires its AHH motif for nuclease activity. Moreover, this toxicity can be specifically neutralized by its cognate immunity protein. While this study was under way, a type VI nuclease effector Tse7 with GHH catalytic site (Tox-GHH2, pfam14412) from the same HNH/ENDO VII superfamily was reported in *P. aeruginosa* (22). In conclusion, HNH nuclease superfamily appeared to be a widespread T6SS nuclease toxin family for antibacterial activity. These results strongly suggested the *v2a-v3a* toxin-immunity relationship, which supported their contribution to 1D1609’s antibacterial activity to *E. coli*.

### VgrG can be exchanged between C58 and 1D1609

Considering that VgrGa, VgrGc and VgrGd of 1D1609 and VgrG2 of C58 are highly similar with identical C-terminal region and their DUF2169 and DUF4150 share high sequence similarity (Table S3), it is plausible that these VgrG proteins can substitute each other’s role in effector delivery. Previous work showed that Hcp and VgrG can be exchanged between *Vibrio cholerae* cells in a T6SS-dependent manner and can be reused to assemble a new T6SS (23). Thus, we hypothesized that *Agrobacterium* may take advantage of having multiple, highly similar VgrG to receive or donate this effector carrier from/to its siblings and the donated VgrG can be used by the recipient cell to kill another competing bacterial cell. To test this hypothesis, we designed a tripartite interbacterial transfer system that will allow us to see the killing of the target *E. coli* cell only when there is a functionally exchangeable VgrG protein translocated from C58 to 1D1609. The nearly identical VgrGs (G2^C58^ and Ga^1D1609^) expressed on a non-transferable pRL662 plasmid in C58 Δ*tdei1*Δ*G1*Δ*G2op*, a mutant deleting *vgrG1, tde1-tdi1, vgrG2* operon, serves as a donor cell. 1D1609 *Δ4* with deletion of all four *vgrG* genes but is still armed with effectors is designated as a recipient cell as well as an attacker to kill *E. coli* (Fig 6A). Since no antibacterial activity can be detected in C58 Δ*tdei1*Δ*G1*Δ*G2op* (14) and 1D1609 *Δ4* (Fig 4B), 1D1609 *Δ4* can only exhibit antibacterial activity by obtaining VgrG proteins from C58 donor and using it for effector delivery to kill *E. coli*.

**Fig 6.**
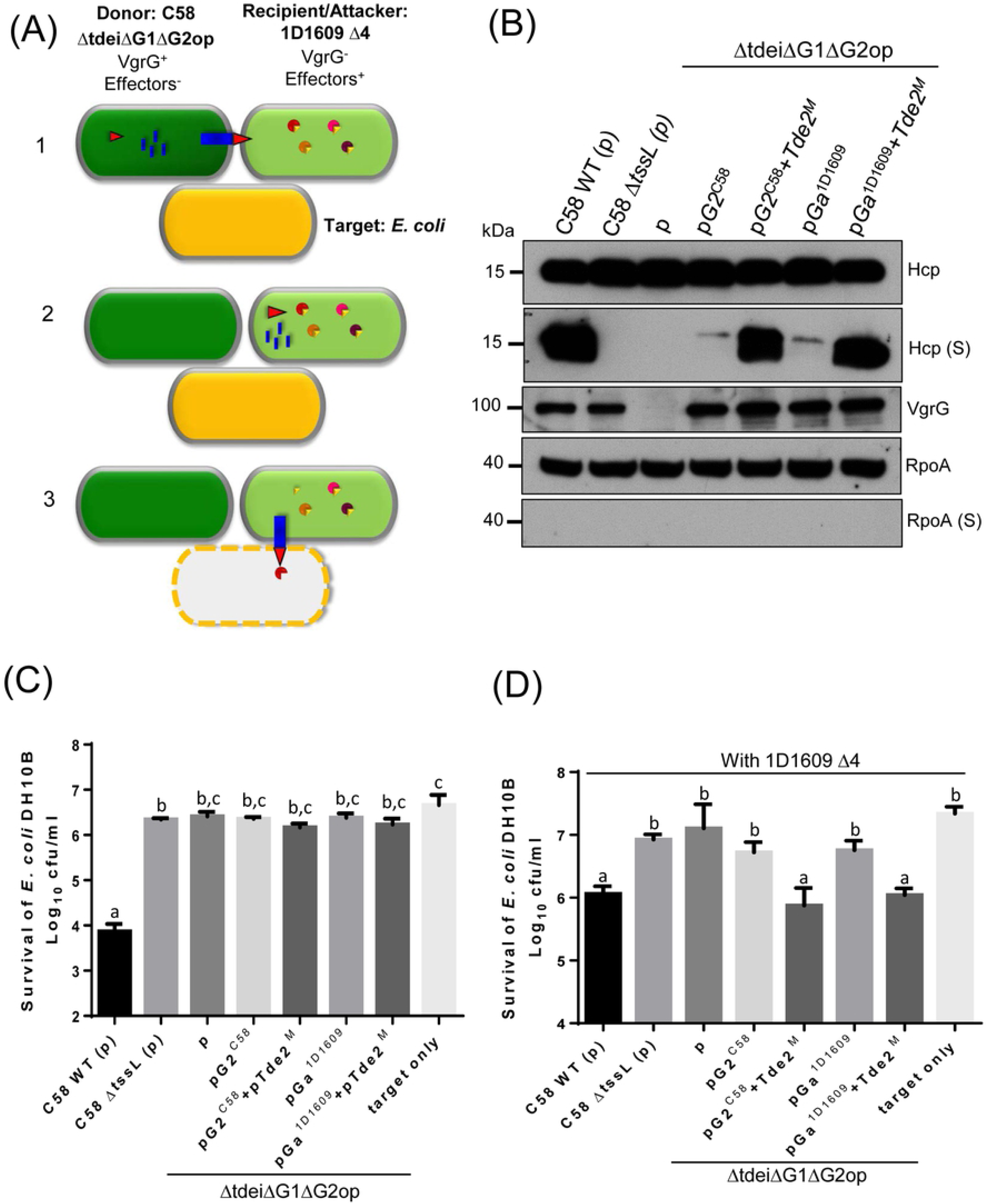
VgrG can be exchanged between C58 and 1D1609 for executing antibacterial activity. **(A)** Schematic diagram for tripartite interaction assay to determine the VgrG exchange/killing of *E. coli* by the recipient *Agrobacterium*. The donor C58 Δ*tdei*Δ*G1*Δ*G2op* was co-cultured with recipient 1D1609 Δ4 and *E. coli* target. The “exchangeable” VgrGs (G2^C58^ or Ga^1D1609^) was expressed alone or co-expressed with Atu3641 and Tde2 with catalytic site mutation (named as Tde2^M^) in donor C58 cells. The exchange/killing is divided in three steps: 1) Donor C58 expresses T6SS components without active toxin effectors and recipient 1D1609 expresses effectors complexed with cognate immunity proteins (EI pairs shown in circle with triangle) without T6SS assembly. T6SS is assembled in donor C58 for injecting Hcp (blue blocks) and VgrG (red triangle) into recipient 1D1609. 2) Recipient 1D1609 uses the translocated VgrG to assemble a new T6SS machine carrying 1D1609 effector. 3) Recipient 1D1609 injects the toxin effector and kills the target *E. coli* **(B)** Hcp secretion in the donor C58. *A. tumefaciens* strain C58 or derivatives containing the plasmid only (p) or expressing the indicated genes were cultured in ABMES (pH 5.5) broth at 25°C and cellular and secreted (S) fractions were subjected to immunoblotting using anti-Hcp, anti-VgrG, and RpoA. The T6SS inactive mutant Δ*tssL* is a negative control and RpoA serves as a non-secreted protein control. **(C)** and **(D)** *A. tumefaciens* 1D1609 strains (indicated in the x axis) was mixed with *E. coli* DH10B cells expressing pRLBla to confer ampicilin resistance at 30:1 ratio in AK medium. The survival of *E. coli* cells was quantified as cfu as shown on the y axis when co-cultured with donor cells only **(C)** or with donor and recipient 1D1609 cells, at 1:10 ratio **(D)**. Data are mean ± SEM of four biological repeats from three independent experiments and letters above the bar indicate significant difference (P < 0.05) determined by Tukey’s HSD test.

During this course of study, our group also discovered that Tde effector loading onto VgrG is critical for T6SS assembly and Hcp secretion (24). Thus, aside from expressing G2^C58^ or Ga^1D1609^ in C58 Δ*tdei*Δ*G1*Δ*G2op* donor strain, a plasmid expressing Tde2 with catalytic site mutation and its cognate DUF2169 chaperone/adaptor (pTde2^M^) was also included to ensure effective T6SS assembly and firing. Hcp secretion and antibacterial activity assays confirmed that the G2^C58^ or Ga^1D1609^ in C58 Δ*tdei*Δ*G1*Δ*G2op* donor strain remained active in Hcp secretion as WT level but completely lost the antibacterial activity against *E. coli* (Fig 6B, C). However, the donor-recipient-prey tripartite co-incubation showed that only the experimental set with C58 Δ*tdei*Δ*G1*Δ*G2op* donor strain harboring G2^C58^ or Ga^1D1609^ in the presence of pTde2^M^ can allow 1D1609 *Δ4* to exhibit antibacterial activity to *E. coli* (Fig 6D). 1D1609 *Δ4* co-cultured with donor strains C58 Δ*tssL*, Δ*tdei*Δ*G1*Δ*G2op* harboring vector only, G2^C58^ or Ga^1D1609^ alone do not exhibit detectable killing activity to *E. coli*. These results suggest that 1D1609 can use the VgrG2^C58^ and VgrGa^1D1609^ delivered from C58 to kill *E. coli* and VgrG2^C58^ can function as an effector carrier in 1D1609.

## Discussion

Functional redundancy is a useful strategy for the pathogen to build a robust system (25). In this study, our phylogenomic analyses revealed that the majority of T6SS-harboring Proteobacteria encode multiple *vgrG* genes. Among them, *A. tumefaciens* strain 1D1609 is unique for having multiple VgrG paralogs (VgrGa, VgrGc, and VgrGd) with high sequence identity (99%). The three *vgrG* genetic modules have the same gene orders encoding highly conserved VgrG- DUF2169-DUF4150. But the DUF4150-linked effectors do not share significant sequence similarities suggesting that these effectors may have distinct functions. This study provided genetic evidence that these near identical VgrG paralogs can carry not only their genetically linked cognate effector but also non-genetically linked effectors sharing the highly similar DUF4150. The functional replacement or exchange can occur for these proteins produced in the same bacterial cells or those taking from its isogenic sibling or close relatives during T6SS attacks. Thus, such functional redundancy may be a beneficial strategy for agrobacteria to compete in their ecological niche.

While several studies clearly demonstrated that VgrG is specifically required for delivery of cognate effector(s) encoded in the same *vgrG* genetic module (7, 8) and C-terminal variable region of VgrG is the molecular determinant (9, 10), only few studies reveal the roles of highly homologous VgrG proteins in effector delivery. *D. dadantii* 3937 also encodes two highly similar VgrG homologs (VgrG_A_ and VgrG_B_, Table S1) which are both genetically linked to genes encoding homologous DUF1795-containing EagR chaperone and Rhs-linked effectors RhsA and RhsB (26). The homologous DUF1795 shares 97% amino acid sequence identity and RhsA and RhsB share 86% amino acid sequence identity with conserved N-terminal PAAR domain (99% identity). By sharing the highly similar chaperone and PAAR-Rhs regions, RhsA and RhsB effectors each harboring distinct C-terminal nuclease toxin domains could be delivered by not only its cognate but also non-cognate VgrG. Indeed, the competition assay revealed that either *vgrG_A_* or *vgrG_B_* is required for RhsB-mediated inhibition. The inhibitor cells without *vgrG_B_*in Δ*rhsA*/*rhsB*^+^ background can still have competitive advantage. The advantage is lost when both *vgrG*s are deleted and restored when *vgrG_B_* was expressed in double *vgrG* deletion mutant. Sharing the same VgrG carrier was also found in *Serratia marcescens* Db10, in which two different PAAR-domain containing Rhs effectors (RhS1 and RhS2) can functionally pair with the same VgrG protein (VgrG2) for delivery to target cells (27). The *rhs2* effector gene is not genetically linked to *vgrG2*, but both Rhs1 and Rhs2 have the same DUF1795 (EagR)-PAAR-Rhs gene organization. The EagR1 share only 26% amino acid sequence identity with EagR2 and each specifically interacts with and required for delivery of its cognate Rhs (27, 28). VgrG2 can form VgrG2-Rhs1-EagR1 or VgrG2-Rhs2-EagR2 complexes even though the N-terminal PAAR and full-length amino acid sequence of Rhs1 and Rhs2 share only 34% and 30% identity, respectively. In this case, high amino acid sequence identity in chaperone and PAAR is not required for functional exchange. Indeed, recent studies revealed However, they may share structural similarity that is sufficient for interacting with the same VgrG.

A recent study demonstrated that T6SS components, Hcp and VgrG, can be transferred and reused by *V. cholera* cells (23). An isogenic LacZ^+^ reporter strain was used to measure the level of LacZ released for detection of cell lysis when the isogenic donor cells translocate Hcp and VgrG2 to isogenic recipient cells for T6SS assembly and firing. In our system, a tripartite interbacterial exchange/competition assay was designed to show evidence that highly similar VgrG proteins can be exchanged between two different *A. tumefaciens* genomospecies to kill the co-existing *E. coli*. Different *A. tumefaciens* genomospecies are known to co-exist in the same geographic area (29). Several studies revealed that more than two genomospecies of *A. tumefaciens* can be found in the same crown gall or soil samples (30, 31). Using tomato and maize seedlings, Gharsa et al., (32) evaluated the competition of different *A. tumefaciens* genomospecies in soil and rhizosphere and concluded that related ecotypes can coexist together. The initial competition alters the relative abundance, but does not eliminate the weaker strain. VgrG exchange may bring the advantages of sympatry and possessing different sets of T6SS gene clusters can maintain balance in sympatry. In *V. cholerae*, the bacterial strains that are incompatible (i.e. with different effector module sets), actively use their T6SS to kill susceptible strain during intra-species competition (33). However, in *A. tumefaciens* species, compatibility of T6SS EI pairs are not always predictive of competition outcomes (15). C58 and 1D1609, which belongs to different genomospecies, can exhibit antibacterial activity against each other when the attacker is initiated with a higher relative abundance during an *in planta* interbacterial competition assay. However, C58 and 1D1609 can coexist with similar competitiveness when co-cultured at equal ratio despite their distinct EI pairs (Fig S4). No significant difference of competitiveness (competitive index ∼1) could be observed when C58 WT is co-inoculated with equal number of 1D1609 WT or *ΔtssL*, or vice versa. This indicates that the two strains are proliferating equally *in planta* regardless of the presence or absence of a functional T6SS in different host plants. Interestingly, highly similar VgrGs are common in different *A. tumefaciens* genomospecies. In genomospecies 1 (G1), all (except 1D1108) share 100% identity while G8 (represented by C58 VgrG1 and two VgrGs in 1D132) share 95.7-99.8%. G7 (three VgrGs in 1D1609) and G8 (C58 VgrG2, 1D132 VgrG and three VgrGs in 1D1609) share 96.9-99.9% identity (Fig S3). Thus, VgrG exchange might be common between different genomospecies, which may allow them to coexist and could in turn benefit *A. tumefaciens* sympatry in its ecological niche.

While not always evident by statistical analysis, deletion of single *vgrGa* (*ΔvgrGa*) or its associated effector-immunity (*ΔEIa*) showed modest reduction of antibacterial activity (Fig 4A, 5A, B), suggesting the primary role of *vgrGa* genetic module in antibacterial activity of 1D1609. The *vgrGa*-associated effector V2a belongs to different type of nuclease toxin distinct from the Tde1 and Tde2 nuclease toxins (toxin_43 domain with HxxD catalytic motif) identified in C58 (14). The use of enzymatic toxins such as nuclease is the most prominent theme in bacterial warfare (25). Using bioinformatics analysis, Ma et al (35) found that among more than 2,500 T6SS-dependent Rhs-CTs with N-terminal PAAR in 143 bacterial species, nuclease represents the major group. Among these, the Rhs-AHH is the most dominant nuclease found in 66 bacterial species. Effectors with predicted AHH nuclease has been reported with antibacterial activity in *V. parahaemolyticus* VP1415 (36) and *Pectobacterium carotovorum subsp. brasiliense* strain PBR 1692 (37) but no nuclease activity has been tested. In *Acinetobacter baylyi*, it was shown that the genomic DNA was completely degraded when Rhs2 with C-terminal AHH motif is expressed in *E. coli* (38). In this study, we characterize further to show that the AHH motif is indeed responsible for the DNase activity (Fig. 5). The other conserved motif of HNH/ENDO VII superfamily nuclease has been reported in *P. putida* KT2440 Tke2 (Rhs-Tox-HNH) and Tke4 (Rhs-Tox-SHH) (39) and functional characterization was done only in Tox-GHH2 in Tse7 in *P. aeruginosa* (*22*). Among the 100 Soft-rot Enterobacteriaceae (SRE) genomes surveyed by Bellieny-Rabelo et al (37), about a quarter encodes AHH, suggesting that this nuclease is a common weapon deployed by both plant and animal pathogenic bacteria in interbacterial competition.

Among the four *vgrG* clusters encoded in 1D1609, *vgrGb* is distinct from the other three. Our data suggest that *vgrGb* and *v4b* encoding a putative toxin does not seem to play a role in antibacterial activity (Fig. 4 and 5A, B). This may be due to low expression level of this gene. Alternatively, V4b may target eukaryotic cells because it contains a nuclear localization signal and has an extensive structural similarity to the insecticidal TcdB2-TccC3 toxin of *Photorhabdus luminescens* and YenC2 toxin of *Yersinia entomophaga* (Table S2). The activity of the remaining effectors remains unknown and is currently under investigation.

In conclusion, *Agrobacterium* has developed a flexible mode of T6SS effector delivery, by using highly similar VgrGs, either produced endogenously or injected from its close relatives, for T6SS assembly and firing. Since VgrG and PAAR are among the least abundant T6SS components (34) and limiting VgrG can result in reduced number of T6SS assemblies (23). having multiple highly similar VgrGs may enhance robustness and abundance of T6SS assemblies. This flexibility could allow them to be an ally to its sister cells and an effective weapon against distantly related bacterial competitors. The knowledge gained in this study can help advance the understanding of mechanisms and physiological roles of multiple VgrG proteins and associated effectors and may provide new insights for sympatric speciation.

## Materials and Methods

### Bacterial Strains, plasmids, and growth conditions

The bacterial strains, plasmids and primers used in this study are listed in Table S4 and S5. *A. tumefaciens* strains were grown in 523 medium at 25 °C while *E. coli* strains were grown in LB medium at 37 °C. In-frame deletion mutants in *Agrobacterium* were generated using a suicide plasmid via double crossing over by electroporation or by conjugation (40). The mutants were confirmed by colony PCR and/or Southern blot analysis (Fig S5). Site directed mutagenesis was done using overlapping PCR (41).

### Phylogenetic and Sequence Analysis

All bioinformatics tools were used with the default parameters unless stated otherwise. The function of putative effectors were inferred from NCBI’s Conserved Domain Database (CDD) based on BLASTP searches (18, 19) and Protein Homology/analogy Recognition Engine (Phyre2) (17). SignalP was used to predict signal peptides (42). PSORTb version 3.0.2 was used to predict sub-cellular localization of proteins (43).

For the phylogenomic analysis to investigate the VgrG diversity among Proteobacteria, the VgrG homologs from five representatives (i.e., *A. tumefaciens* 1D1609, *Burkholderia thailandensis* E264, *Geobacter sulfurreducens* PCA, *Helicobacter cinaedi* CCUG 18818, and *P. aeruginosa* PAO1) were used as queries to run BLASTP searches against the NCBI non-redundant protein database (e-value cutoff = 1e-15, max target sequences = 100,000). Spurious hits, defined as hits with the high scoring pairs (HSP) accounting for <74% of the query sequence or sequence similarity <48% within the HSP, were removed. The resulting lists from different query species were combined to remove redundant hits. The combined list was further manually curated to keep only the targets with complete genome sequences available in GenBank. Based on the taxonomy information of the targets, the list was iteratively trimmed to achieve a balance taxon sampling with ∼50 genomes to represent all major lineages within Proteobacteria. One Planctomycetes, *Pirellula staleyi* DSM 6068, was included as the outgroup to root the species phylogeny (Fig 1). In addition to the aforementioned data set for phylum-level (i.e., Proteobacteria) diversity, a second data set containing 11 *Agrobacterium* genomes (15) was compiled to examine the genus-level VgrG diversity; *Neorhizobium galegae* HAMBI 540 was included as the outgroup for this second data set (Fig. S3).

After the genome selection, the procedures for homologous gene identification and molecular phylogenetic analysis were based on those described in our previous studies (44, 45). Briefly, the homologous genes among all genomes were identified by using OrthoMCL (46). The single-copy genes shared by all genomes were used for inferring the species phylogeny. Additionally, all *vgrG* homologs in these genomes, defined as the genes containing at least 80% of the VgrG domain (TIGR03361), were extracted for inferring the gene tree. The protein sequences were aligned using MUSCLE (47) and the molecular phylogenies were inferred using MrBayes (48). The amino acid substitution model was set to mix with gamma-distributed rate variation across sites and a proportion of invariable sites. The number of rate categories for the gamma distribution was set to four. The Markov chain Monte Carlo analysis was set to run for 1,000,000 generations and sampled every 100 generations. The first 25% of the samples were discarded as the burn-in. Furthermore, the protein sequence identity among all VgrG homologs were calculated using PROTDIST (49). The amino acid sequence identity and similarity percentage of VgrG and VgrG-associated proteins were determined using Vector NTI Advance 11.5.4.

### Secretion Assay

Secretion assay from liquid culture of *A. tumefaciens* grown in either LB rich medium or AB-MES minimal medium was done as previously described (40). Proteins from cellular and secreted fractions were resolved by SDS-PAGE and transferred onto a PVDF membrane by using a transfer apparatus (Bio-Rad). The membrane was probed with primary antibody against C58 VgrG1 (1:1,000), which recognizes the C58 VgrG2 and all four VgrGs of 1D1609, Hcp (1:2,500), and RpoA (1:7,500) (20), followed by incubation with horseradish peroxidase-conjugated anti-rabbit secondary antibody (1:30,000) and visualized with the ECL system (Perkin**-**Elmer).

### Interbacterial Competition Assay

*E.* co*li* killing assay was performed as described previously (9). In brief, overnight culture of *A. tumefaciens* and *E. coli* strain were adjusted to OD_600_ 0.1 and incubated at 25 °C for 4 h. *A. tumefaciens* and *E. coli* cells were mixed at a 30:1 ratio (OD 0.3:0.01) and spotted on LB agar (1.5%) plates. After 18 hr incubation at 25 °C, the co-cultured cells were collected, serially diluted and plated on LB agar plate containing appropriate antibiotics to quantify surviving *E. coli* by counting colony forming unit (cfu). When enhanced killing activity is desired, an optimized growth medium, named as *<u>A</u>grobacterium* <u>K</u>ill-triggering, AK medium (3 g K_2_HPO_4_, 1 g NaH_2_PO_4_, 1 g NH_4_Cl, 0.15 g KCl, 9.76 g MES in 900 mL ddH20, pH5.5) modified from AB-MES medium and solidified by 2% (w/v) agar, was used instead. Data are expressed as mean ± SEM (standard error of the mean) from three independent experiments. Statistics was done using one-way ANOVA and Tukey’s honestly significance difference (HSD) test (http://astatsa.com/OneWay_Anova_with_TukeyHSD/).

*In planta* bacterial competition assay was performed as described previously (14). *A. tumefaciens* strain C58 transformed with gentamycin resistance-conferring pRL662 plasmid and strain 1D1609 conferring spectinomycin resistance were mixed at 1:1 (OD 1) ratio and infiltrated into 6- to 7-wk-old *Nicotiana benthamiana* and *Medicago truncatula* leaves. The competition outcome was quantified after 24 hours by counting cfu on LB plates with appropriate antibiotics. 1D1609/C58 ratio at t=24 h was divided by 1D1609/C58 ratio at t=0 h to calculate the competitive index.

### Growth Inhibition Assay

Growth inhibition assay was performed as described previously (14). In brief, overnight cultures of *E*. *coli* DH10B strain with vectors or their derivatives were adjusted to OD_600_ 0.1. The expression of the tested immunity protein was induced by 1 mM IPTG for 1 hr before L-arabinose (0.2% final concentration) was added to induce expression of the toxin.

### Plasmid DNA Degradation Analysis in *E. coli* Cells

*In vivo* plasmid DNA degradation analysis was performed as described previously (14). In brief, overnight cultures of *E*. *coli* DH10B strain with empty vectors or derivatives expressing *v2a* and catalytic site mutant were harvested and adjusted to an OD_600_ 0.3. L-arabinose (0.2%) was added to induce the expression and cultured for 2 hr. Equal amounts of cells were used for plasmid DNA extraction and equal volume of extracted DNA was resolved in agarose gel followed by ethidium bromide staining.

### Southern blot hybridization

Southern blot hybridization was performed as described (50). In brief, genomic DNA (gDNA) was extracted from overnight cultures of selected strains using Genomic DNA purification Kit (Promega) as per manufacturer’s instructions. About 35 μg of gDNA was digested using NEB restriction enzymes (Nsil and PvuII) and resolved in 0.8% agarose gel, 50V, 3hr. The DNA fragments were transferred to a positively charged nylon membrane (Roche Diagnostics). Nylon membranes were cross-linked and used for hybridization with DIG-labeled probe (Roche). Hybridization was done overnight at 65°C using hybridization solution (FastHyb-Hybridization solution, BioChain). The washing, blocking and detection were done using Roche Wash and Block Buffer Set and DIG DNA Labeling and Detection Kit, according to manufacturer’s instructions (Roche). The membrane was exposed to X-ray for detection. The probe for the four *vgrGs* (*vgrGa*, *vgrGb*, *vgrGc* and *vgrGd*) and putative toxin genes (*v2a*, *v2c*, *v2d* and *v4b*) was prepared using the plasmid DNA *vgrGa*-pJN105, *v2d*-pJN105 and *v4b*-pJN105, respectively. The primer sets used are listed in Table S5.

## Acknowledgments

We thank members of Lai lab for their help, encouragement and stimulating discussion. We are grateful for Genomic Technology Core of Institute of Plant and Microbial Biology for DNA sequencing.

## Author Contributions

Conceptualization, E.-M.L. and M.-M.S; Data curation, M.-M.S, S.-T.C. and C.-J.C; Methodology, E.-M.L., C.-H.K., C.-F.W., and M.-M.S; Investigation, C.-H.K., S.-T.C., M.-M.S., and C.-J.C.; Project administration, E.-M.L; Resources, E.-M.L; Supervision, E.-M.L. and C.-H.K.; Writing, M.-M.S., C.-H.K., and E.-M.L.; All authors reviewed and approved the manuscript.

## Supporting information captions

**S1 Fig.** Bayesian phylogeny of the VgrG homologs identified in the representative Proteobacteria genomes presented in Fig 1.

The color coding is based on the class: Alphaproteobacteria (black), Betaproteobacteria (blue), Deltaproteobacteria (cyan), Epsilonproteobacteria (purple) and Gammaproteobacteria (green). Numbers on the branches indicate the support levels based on posterior probabilities, only values >60% were shown.

**S2 Fig.** Multiple sequence alignment of VgrG amino acid sequences of 1D1609 and C58. All VgrG homologs were aligned and conserved amino acid residues are shaded in yellow and variable residues are in blue or green, while the C-terminal extension of C58 VgrG1 not homologous to the other VgrGs is unshaded. The red and blue lines indicate the predicted gp27 and gp5 domains, respectively. The amino acid residue number is indicated.

**S3 Fig.** VgrG tree of the 11 *A. tumefaciens* strains. All VgrG homologs in these 11 genomes were extracted for inferring the gene tree. The VgrG orthologs share ≥ 95% identity in amino acid sequence were highlighted. The genomospecies and strain name for each VgrG indicated by a locus tag are shown.

**S4 Fig.** *In planta* competition experiments between C58 and 1D1609 at 1:1 ratio. *A. tumefaciens* strain C58 harboring gentamycin resistance-conferring pRL662 and strain 1D1609 which was selected in spectinomycin plate were mixed at 1:1 ratio and infiltrated into leaves of 6- to 7-wk-old *Nicotiana benthamiana* **(A)** and *Medicago truncatula* **(B)**. The competition outcome was quantified at 0 and 24 hours post-infection by counting cfu on LB plates with appropriate antibiotics. Data are mean ± SEM, with **e**ach data point indicating the competitive index of four to six biological replicates from a total of two to three independent experiments. No statistical difference (P > 0.05) could be detected among different samples as determined by Tukey’s HSD test.

**S5 Fig.** Southern blot analysis of mutants generated in this study. Schematic diagram showing probes used for Southern blotting, restriction enzyme cleavage sites and expected sizes. The genomic DNA is hybridized with **(A)** 821-bp *vgrG* probe with homology to all four *vgrG* genes (*vgrGabcd)* was used to confirm *vgrG* mutants; **(B)** 478-bp *v2* probe with homology to *v2acd* was used to confirm *v2a*, *v2c* and *v2d* mutant and **(C)** 778-bp *v4b* probe was used to confirm *v4b* mutant. Each lane contains 30 µg of genomic DNA digested with NsiI only or combined with PvuII. The expected size (in bp) is indicated on the side of the blot.

**Table S1.** Percentage identity of VgrG within genomes with highly identical multiple VgrGs. The locus tag of each homolog is shown in rows and column and highlight shows ≥ 95% identity.

**Table S2.** Predicted functions/domains of toxin/s associated with each *vgrG* based on Phyre or NCBI CDD search and BLASTP.

**Table S3.** Amino acid sequence similarity/identity among the VgrG loci homologs in C58 and 1D1609

**Table S4.** Bacterial strains and plasmids.

**Table S5.** Primers used in this study.

## References

1. Bingle LE, Bailey CM, Pallen MJ. Type VI secretion: a beginner’s guide. Current opinion in microbiology. 2008;11(1):3–8.

2. Russell AB, Peterson SB, Mougous JD. Type VI secretion system effectors: poisons with a purpose. Nat Rev Micro. 2014;12(2):137–48.

3. Tang JY, Bullen NP, Ahmad S, Whitney JC. Diverse NADase effector families mediate interbacterial antagonism via the type VI secretion system. The Journal of biological chemistry. 2018;293(5):1504–14.

4. Ting S-Y, Bosch DE, Mangiameli SM, Radey MC, Huang S, Park Y-J, et al. Bifunctional Immunity Proteins Protect Bacteria against FtsZ-Targeting ADP-Ribosylating Toxins. Cell. 2018.

5. Zoued A, Brunet YR, Durand E, Aschtgen M-S, Logger L, Douzi B, et al. Architecture and assembly of the Type VI secretion system. Biochimica et Biophysica Acta (BBA) - Molecular Cell Research. 2014;1843(8):1664–73.

6. Leiman PG, Basler M, Ramagopal UA, Bonanno JB, Sauder JM, Pukatzki S, et al. Type VI secretion apparatus and phage tail-associated protein complexes share a common evolutionary origin. Proceedings of the National Academy of Sciences of the United States of America. 2009;106(11):4154–9.

7. Whitney JC, Beck CM, Goo YA, Russell AB, Harding BN, De Leon JA, et al. Genetically distinct pathways guide effector export through the type VI secretion system. Mol Microbiol. 2014;92(3):529–42.

8. Hachani A, Allsopp LP, Oduko Y, Filloux A. The VgrG Proteins Are “à la Carte” Delivery Systems for Bacterial Type VI Effectors. Journal of Biological Chemistry. 2014;289(25):17872–84.

9. Bondage DD, Lin J-S, Ma L-S, Kuo C-H, Lai E-M. VgrG C terminus confers the type VI effector transport specificity and is required for binding with PAAR and adaptor–effector complex. Proceedings of the National Academy of Sciences. 2016;113(27):E3931–E40.

10. Flaugnatti N, Le TT, Canaan S, Aschtgen MS, Nguyen VS, Blangy S, et al. A phospholipase A1 antibacterial Type VI secretion effector interacts directly with the C-terminal domain of the VgrG spike protein for delivery. Mol Microbiol. 2016;99(6):1099–118.

11. Lien YW, Lai EM. Type VI Secretion Effectors: Methodologies and Biology. Frontiers in cellular and infection microbiology. 2017;7:254.

12. Gelvin SB. AGROBACTERIUM AND PLANT GENES INVOLVED IN T-DNA TRANSFER AND INTEGRATION. Annual review of plant physiology and plant molecular biology. 2000;51:223–56.

13. Hwang H-H, Yu M, Lai E-M. Agrobacterium-Mediated Plant Transformation: Biology and Applications. The Arabidopsis Book. 2017:e0186.

14. Ma L-S, Hachani A, Lin J-S, Filloux A, Lai E-M. *Agrobacterium tumefaciens* Deploys a Superfamily of Type VI Secretion DNase Effectors as Weapons for Interbacterial Competition In Planta. Cell Host & Microbe. 2014;16(1):94–104.

15. Wu C-F, Santos MNM, Cho S-T, Chang H-H, Tsai Y-M, Smith DA, et al. Plant pathogenic Agrobacterium tumefaciens strains have diverse type VI effector-immunity pairs and vary in in planta competitiveness. Molecular Plant-Microbe Interactions. 2019.

16. Bladergroen MR, Badelt K, Spaink HP. Infection-Blocking Genes of a Symbiotic Rhizobium leguminosarum Strain That Are Involved in Temperature-Dependent Protein Secretion. Molecular Plant-Microbe Interactions. 2003;16(1):53–64.

17. Kelley LA, Mezulis S, Yates CM, Wass MN, Sternberg MJ. The Phyre2 web portal for protein modeling, prediction and analysis. Nature protocols. 2015;10(6):845–58.

18. Marchler-Bauer A, Bo Y, Han L, He J, Lanczycki CJ, Lu S, et al. CDD/SPARCLE: functional classification of proteins via subfamily domain architectures. Nucleic Acids Research. 2017;45(Database issue):D200–D3.

19. Boratyn GM, Camacho C, Cooper PS, Coulouris G, Fong A, Ma N, et al. BLAST: a more efficient report with usability improvements. Nucleic Acids Research. 2013;41(Web Server issue):W29–W33.

20. Lin J-S, Ma L-S, Lai E-M. Systematic Dissection of the Agrobacterium Type VI Secretion System Reveals Machinery and Secreted Components for Subcomplex Formation. PLOS ONE. 2013;8(7):e67647.

21. Zhang D, Iyer LM, Aravind L. A novel immunity system for bacterial nucleic acid degrading toxins and its recruitment in various eukaryotic and DNA viral systems. Nucleic Acids Research. 2011;39(11):4532–52.

22. Pissaridou P, Allsopp LP, Wettstadt S, Howard SA, Mavridou DAI, Filloux A. The *Pseudomonas aeruginosa* T6SS-VgrG1b spike is topped by a PAAR protein eliciting DNA damage to bacterial competitors. Proceedings of the National Academy of Sciences. 2018;115(49):12519.

23. Vettiger A, Basler M. Type VI Secretion System Substrates Are Transferred and Reused among Sister Cells. Cell. 2016;167(1):99–110.e12.

24. Lien Y-W, Wu C-F, Bondage D, Lin J-S, Shih Y-L, H. Chang J, et al. Effector loading onto VgrG spike proteins is critical for the assembly of the type VI secretion system in Agrobacterium tumefaciens. bioRxiv preprint first posted online Feb. 21, 2019; doi: http://dx.doi.org/10.1101/556712.

25. Galán JE. Common themes in the design and function of bacterial effectors. Cell host & microbe. 2009;5(6):571–9.

26. Koskiniemi S, Lamoureux JG, Nikolakakis KC, t’Kint de Roodenbeke C, Kaplan MD, Low DA, et al. Rhs proteins from diverse bacteria mediate intercellular competition. Proceedings of the National Academy of Sciences of the United States of America. 2013;110(17):7032–7.

27. Cianfanelli FR, Alcoforado Diniz J, Guo M, De Cesare V, Trost M, Coulthurst SJ. VgrG and PAAR Proteins Define Distinct Versions of a Functional Type VI Secretion System. PLOS Pathogens. 2016;12(6):e1005735.

28. Alcoforado Diniz J, Coulthurst SJ. Intraspecies Competition in Serratia marcescens Is Mediated by Type VI-Secreted Rhs Effectors and a Conserved Effector-Associated Accessory Protein. Journal of Bacteriology. 2015;197(14):2350–60.

29. Vogel J, Normand P, Thioulouse J, Nesme X, Grundmann GL. Relationship between spatial and genetic distance in Agrobacterium spp. in 1 cubic centimeter of soil. Applied and environmental microbiology. 2003;69(3):1482–7.

30. Costechareyre D, Rhouma A, Lavire C, Portier P, Chapulliot D, Bertolla F, et al. Rapid and Efficient Identification of Agrobacterium Species by recA Allele Analysis. Microbial Ecology. 2010;60(4):862–72.

31. Bouri M, Chattaoui M, Gharsa HB, McClean A, Kluepfel D, Nesme X, et al. Analysis of Agrobacterium populations isolated from tunisian soils: genetic structure, avirulent-virulent ratios and characterization of tumorigenic strains. Journal of Plant Pathology. 2016;98(2):265–74.

32. Ben Gharsa H, Bouri M, Glick B, Gannar A, Mougou Hamdane A, Rhouma A. Evaluation of the interspecific competition within Agrobacterium spp. in the soil and rhizosphere of tomato (Solanum lycopersicum) and maize (Zea mays)2018.

33. Unterweger D, Miyata ST, Bachmann V, Brooks TM, Mullins T, Kostiuk B, et al. The Vibrio cholerae type VI secretion system employs diverse effector modules for intraspecific competition. Nature communications. 2014;5:3549.

34. Lin L, Lezan E, Schmidt A, Basler M. Abundance of bacterial Type VI secretion system components measured by targeted proteomics. Nature communications. 2019;10(1):2584.

35. Ma J, Sun M, Dong W, Pan Z, Lu C, Yao H. PAAR-Rhs proteins harbor various C-terminal toxins to diversify the antibacterial pathways of type VI secretion systems. Environmental Microbiology. 2017;19(1):345–60.

36. Salomon D, Kinch LN, Trudgian DC, Guo X, Klimko JA, Grishin NV, et al. Marker for type VI secretion system effectors. Proceedings of the National Academy of Sciences. 2014;111(25):9271–6.

37. Bellieny-Rabelo D, Tanui CK, Miguel N, Kwenda S, Shyntum DY, Moleleki LN. Transcriptome and Comparative Genomics Analyses Reveal New Functional Insights on Key Determinants of Pathogenesis and Interbacterial Competition in *Pectobacterium* and *Dickeya* spp. Applied and environmental microbiology. 2019;85(2):e02050–18.

38. Fitzsimons TC, Lewis JM, Wright A, Kleifeld O, Schittenhelm RB, Powell D, et al. Identification of Novel Acinetobacter baumannii Type VI Secretion System Antibacterial Effector and Immunity Pairs. Infection and Immunity. 2018;86(8):e00297–18.

39. Bernal P, Allsopp LP, Filloux A, Llamas MA. The Pseudomonas putida T6SS is a plant warden against phytopathogens. ISME J. 2017;11(4):972–87.

40. Ma LS, Lin JS, Lai EM. An IcmF family protein, ImpLM, is an integral inner membrane protein interacting with ImpKL, and its walker a motif is required for type VI secretion system-mediated Hcp secretion in Agrobacterium tumefaciens. J Bacteriol. 2009;191(13):4316–29.

41. Ho SN, Hunt HD, Horton RM, Pullen JK, Pease LR. Site-directed mutagenesis by overlap extension using the polymerase chain reaction. Gene. 1989;77(1):51–9.

42. Petersen TN, Brunak S, von Heijne G, Nielsen H. SignalP 4.0: discriminating signal peptides from transmembrane regions. Nat Meth. 2011;8(10):785–6.

43. Yu NY, Wagner JR, Laird MR, Melli G, Rey S, Lo R, et al. PSORTb 3.0: improved protein subcellular localization prediction with refined localization subcategories and predictive capabilities for all prokaryotes. Bioinformatics (Oxford, England). 2010;26(13):1608–15.

44. Lo W-S, Chen L-L, Chung W-C, Gasparich GE, Kuo C-H. Comparative genome analysis of Spiroplasma melliferumIPMB4A, a honeybee-associated bacterium. BMC Genomics. 2013;14(1):22.

45. Lo W-S, Kuo C-H. Horizontal Acquisition and Transcriptional Integration of Novel Genes in Mosquito-Associated Spiroplasma. Genome Biology and Evolution. 2017;9(12):3246–59.

46. Li L, Stoeckert CJ, Roos DS. OrthoMCL: Identification of ortholog groups for eukaryotic genomes. Genome Res. 2003;13.

47. Edgar RC. MUSCLE: multiple sequence alignment with high accuracy and high throughput. Nucl Acids Res. 2004;32.

48. Ronquist F, Huelsenbeck JP. MrBayes 3: Bayesian phylogenetic inference under mixed models. Bioinformatics (Oxford, England). 2003;19(12):1572–4.

49. Felsenstein J. PHYLIP - Phylogeny Inference Package (Version 3.2). Cladistics. 1989;5.

50. Sambrook J. Molecular cloning : a laboratory manual: Third edition. Cold Spring Harbor, N.Y. : Cold Spring Harbor Laboratory Press, [2001] ©2001; 2001.

